# Ctenophore mesoglea is composed of mucins and novel polysaccharides

**DOI:** 10.64898/2026.02.02.702917

**Authors:** Sara Siwiecki, Stephanie Archer-Hartmann, Ian Black, Ikenna Ndukwe, Jiri Vlach, Parastoo Azadi, Casey Dunn, Alison Sweeney

## Abstract

Ctenophora are one of the earliest, if not the earliest branching animal, making them crucial for insight into the early events of metazoan evolution. Ctenophores are largely composed of an extracellular mesoglea, but they lack homologs for the fibrous collagens that form a typical metazoan extracellular matrix (ECM). Therefore, the nature of this body-dominating material and its physiological role remains unknown. We found that ctenophore mesoglea is dominated by mucin-related proteins and novel sulfated polysaccharides. Mucus-like material that functions as an internal skeleton rather than as an external secretion indicates fundamental differences in epithelial function and ECM physiology between ctenophores and other Metazoa, suggesting an early period of biochemical and biomechanical diversity prior to the innovation of elastic connective tissue.

## Main Text

Ctenophores are one of the earliest extant (possibly the earliest) branching animal taxa (Fig. 1A) and live in nearly every environment of the world’s oceans, across wide temperature, pressure, and salinity ranges (*1–5*). Despite their key position for understanding animal evolution and ubiquity in marine environments, many fundamental aspects of their physiology are unexplored.

**Figure 1:**
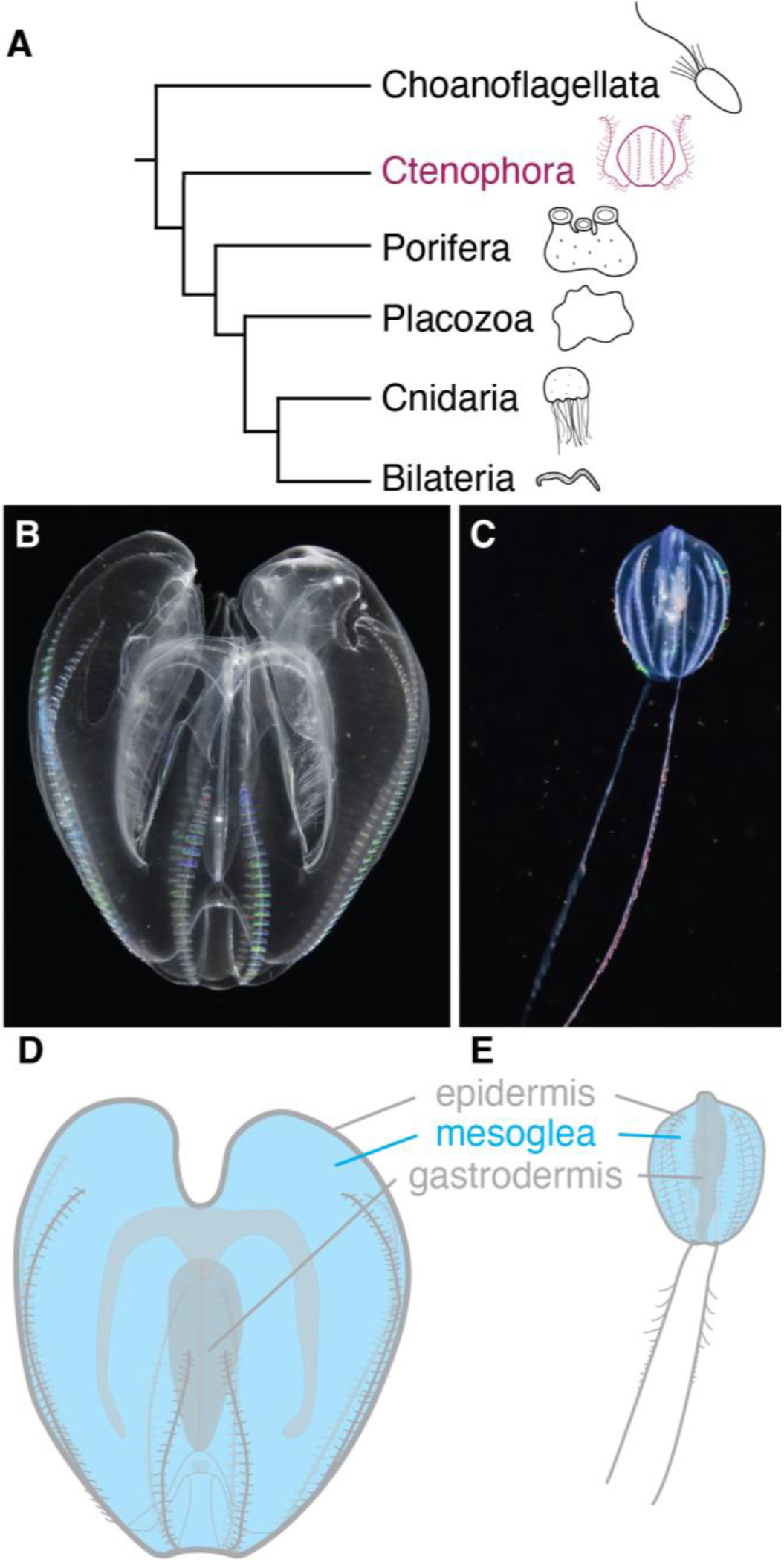
Ctenophores have bulk gelatinous mesogleal tissue. (**A**) Ctenophores are one of our most distant animal relatives. (**B**) *Mnemiopsis leidyi* is a lobate ctenophore, photo taken by SERC/MarineGEO Chesapeake BioBlitz Team. (**C**) *Pleurobrachia pileus* is a cydippid ctenophore. Image cropped from ”Sea Gooseberry (Pleurobrachia pileus) in Norway” by Vsevolod Rudyi, licensed under CC BY 4.0. (**D and E**) The simplified anatomy of *M. leidyi* and *P. pileus*, respectively, comprising gelatinous mesogleal tissue between an outer epidermal layer and an inner gastrodermal layer.

The ctenophore body consists of two thin epithelial layers, the epidermis and the gastrodermis, separated by a voluminous layer of gelatinous mesoglea, illustrated in Fig. 1C and 1D (*6–8*). The mesoglea contains sparse, isolated cells interspersed with bulk extracellular hydrogel material of largely unknown composition. This gelatinous mesoglea forms ∼99% of the organism’s volume, yet few studies have reported on its composition or physiological role. Franc (1985) provided the first, and only, focused description of the biochemical composition of the mesoglea. Franc hypothesized that the mesoglea is formed by a loose three-dimensional network of glycoproteins and sulfated polysaccharides (*7*). This study and others have also observed sparse ”bundle” structures containing polymers with superficial similarity to collagens (*7, 9, 10*). However, ctenophores lack the genes coding for most collagens, and specifically lack homologs to the fibrous, gel-forming collagens that make up the volume-spanning extracellular tissues of other animals (*11, 12*). Ctenophores do have homologs to type IV collagen, which contributes to the sheet-like extracellular basement membrane of metazoan epithelia rather than expanded extracellular tissues (*11–13*). Similarly, ctenophores appear to lack fibronectin or tenascin, further differentiating the expanded extracellular material of ctenophores from other metazoans (*12*). Imaging studies, such as those from Norekian & Moroz (2018), have revealed that ctenophore mesoglea supports neuromuscular fibers that extend inward from epithelial layers (*14, 15*). Because the mesoglea dominates the body volume, prior compositional measurements of whole animals presumably reveal some information about its composition. Ctenophores’ organic material has been previously assumed to be mostly protein, with little lipid or carbohydrate (*16–24*). Most lipids in ctenophores are membrane phospholipids, so are unlikely to participate in the mesoglea structure (*25*). However, elemental analyses show significant non-protein nitrogen, suggesting the presence of uncharacterized glycans, though these have not been carefully assayed (*18*).

Despite ctenophores’ key evolutionary position, there has been no modern description of the body-dominating mesoglea. Thus, we present the first thorough biochemical characterization of ctenophore mesoglea. We focus on *Mnemiopsis leidyi* and *Pleurobrachia pileus*, two species present in the Long Island Sound that represent different clades within ctenophores, Nuda and Tentaculata, respectively (Fig. 1B,C).

Unexpectedly, we found that the mesoglea appears more similar to metazoan mucus than the fibrous-collagen-dominated extracellular matrices (ECMs) of other metazoans. For this study, we define mucus to be an extracellular aqueous solution rich in mucin proteins and complex polysaccharides. Since animal mucuses are typically secreted to epithelial surfaces that are topologically external to the organism, such as skin and gut linings, this finding is surprising and potentially marks a novel mucus function as a volume-filling skeleton. We also report a surprising lack of evidence for covalent linkage between novel mesoglea-constituent sulfated polysaccharides and the mucin protein homologs present in mesoglea, which if confirmed with more specialized experiments, would mark a major chemical structural difference between ctenophore mesogleal mucins and other metazoan mucins.

## Ctenophore mesoglea contains mucin homologs

We identified extracellular proteins in mesoglea using MS-based proteomics with predicted proteins from the chromosome-level northern Atlantic *M. leidyi* genome and the *H. californensis* genome for *P. pileus* (*4, 26*). For each MS hit, we used BLASTp to annotate predicted biological function based on sequence identity with known proteins. Of the extracellular proteins identified, mucus-related proteins were the most represented family (Fig. 2A,B). The mucin domain has high polymorphism (*27*), so protein domain architecture must also be analyzed to determine possible mucin homology. The consensus definition of a functional gel-forming mucin is the serial arrangement of several protein domains: repeated VWD-C8-TIL domains with a C-terminal PTS domain (*27*). We analyzed proteins in our dataset with mucin-like homology using InterProScan and PTSPred search (*28, 29*). While all predicted mucin-like transcripts in these ctenophores had at least a partial set of these domains, we located the full arrangement in one set of proteins in a tandem array in the genome of both species (Fig. 2C). For *M. leidyi*, evm.model.c7.706 contains a partial VWD-C8-TIL repeat and is in a tandem array with evm.model.c7.707, which contains the remaining parts of the repeat and a short PTS domain. Interestingly, these two tandem genes are in a head-to-head orientation with gene evm.model.c7.705, which also contains all the necessary mucin components; thus, if expressed together, these genes may build a larger mucin protein complex. This possibility was also noted by Lang and colleagues (2016), who identified mucin proteins in both *M. leidyi* and *P. bachei*. For *P. pileus*, we identified an MS hit (Hcv1.av93.c7.g869.i1) using the *H. californensis* genome that contains all the mucin components in the definitional arrangement but with a very short PTS domain. Similar to *M. leidyi*, a nearby gene, Hcv1.av93.c7.g874.i1, on the opposite strand has multiple PTS domains. A tandem gene immediately upstream also contains all mucin-related domains in definitional arrangement, Hcv1.av93.c7.g868.i1. If expressed together, these regions potentially encode two mucin proteins. While we do not find all of these proteins in our proteomics results, we do find at least one of them expressed in both species.

**Figure 2:**
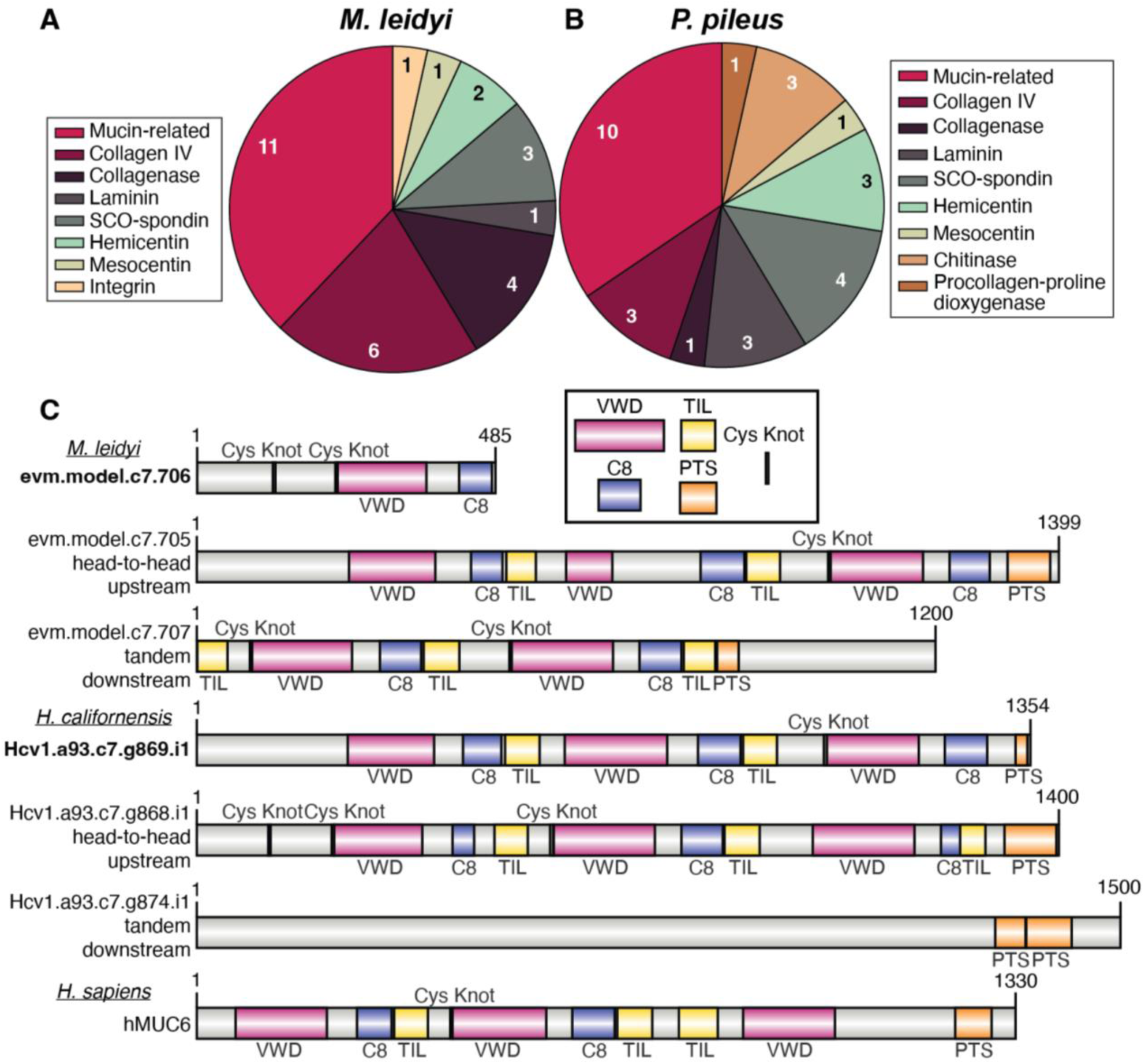
Mucin-related proteins dominate mesoglea protein content. (**A**) The major extracellular proteins identified in *M. leidyi* from LC-MS/MS proteomics. (**B**) The major extracellular proteins identified in *P. pileus* from LC-MS/MS proteomics using an *H. californensis* database. (**C**) Domain architecture of *M. leidyi* and *H. californensis* proteins with homology to metazoan mucins. Proteins that were observed in the proteomics results are noted in bold, and proteins with mucin-related domains that are either in tandem array or head-to-head orientation with the expressed proteins are also shown. hMUC6 is a human mucin protein with the characteristic organization of mucin functional domains. *Homo sapiens* mucin-6 isoform X3 was used (NCBI: XP 054185958.1).

The VWD, TIL, C8, plus PTS domain architecture is a conserved feature of metazoan mucins, but the complete functions of these domains are not yet known, so it is difficult to extrapolate insights to ctenophore mucin structure and function. One motif with established function in mucins is the use of cysteine-rich domains CysD, cysteine knots (CK), and C8, for cross-linking polymerization via disulfide bonds (*30–35*). In human mucins, CysD domains are interspersed within the PTS regions and act as additional locations for interstrand disulfide bonding (*31*). In contrast, we did not identify any CysD domains in the PTS regions of the ctenophore mucins (Fig. 2C). However, a few human mucins, such as MUC19 and MUC6, also lack CysD domains within the PTS region, so this domain is not an absolute requirement for polymerization (*30*). Interestingly, the presence of multiple CysD domains increases the stiffness of mucins and mucus-related molecules (*32*), so the absence of CysD in ctenophore mucins is possibly related to the liquid-like quality of ctenophore mesoglea. Similarly, CK motifs play roles in C-terminal dimerization, the first step of the mucin assembly process (*33*). The C-terminus of ctenophore mucins lacks the CK motif, though there are multiple CK sites throughout the VWD-TIL-C8 repeats along the N-terminus (separate from the C8 domains). These N-terminal cysteines may also contribute to dimerization, both via disulfide bonds and hydrophobic interactions (*31, 35*). We infer from these observations that ctenophore mucins are likely capable of further polymerization, but the differences in the regions responsible for multimerization are large enough that the resulting assemblies are likely to be quite structurally different from other metazoan mucin complexes.

The other mucin-related proteins we identified (Fig. 2A,B) do not all have typical mucin domains, but may complex with other proteins to perform other mucus-related functions. For example, mucus often has a high concentration of actins, myosins, and other cytoskeletal proteins, which play both structural and immune-related roles in these materials (*36, 37*). We similarly identified numerous actins, myosins, and tubulins in ctenophore mesoglea (data S1–S4), which may participate in unknown internal mucus functions or could also be debris of sheared neuromuscular fibers (*14, 15*). However, the relative abundance of these cytoskeletal proteins makes it seem unlikely that cellular debris is the sole source of these proteins in our samples. We also identified many immune-related proteins (data S1 and S3) that often act in concert with mucin proteins (*36, 38*). Significant further research will be required to understand the functions of these proteins in the ctenophores’ internal milieu.

## Novel glycans in ctenophore mesoglea

Glycoproteins, proteoglycans, glycosylaminoglycans (GAGs), and free polysaccharides are constituent components of characterized metazoan connective tissue and mesoglea (*18, 39–46*). We searched for these macromolecules in mesoglea via Nuclear Magnetic Resonance (NMR) Spectroscopy. Various NMR experiments were performed with two protease-treated *M. leidyi* mesoglea biological replicates; spectra that produced solvable structures are presented here. In Figure 3A, a representative ^1^H-^13^C HSQC NMR spectrum for one of the *M. leidyi* samples demonstrates two sulfated-galactose-containing polymers (other replicates and DEAE fractionation confirm residues shown in fig. S1A,B; major peaks are described in table S1). The two solved structures are shown in Figure 3B: a sulfated galactan (residues A and B) and a sulfated Gal-Glc disaccharide (residues C and D) bound to a hydroxylysine residue (Hyl, residue E). These structures are consistent with our findings of high galactose content in gas chromatography-mass spectroscopy (GC-MS) (chromatograms shown in fig. S2A, quantified in table S2). Gal-Glc-Hyl without sulfation is common in collagens (*47–50*), but to our knowledge a sulfated Gal-Glc-Hyl residue is a novel structure.

**Figure 3:**
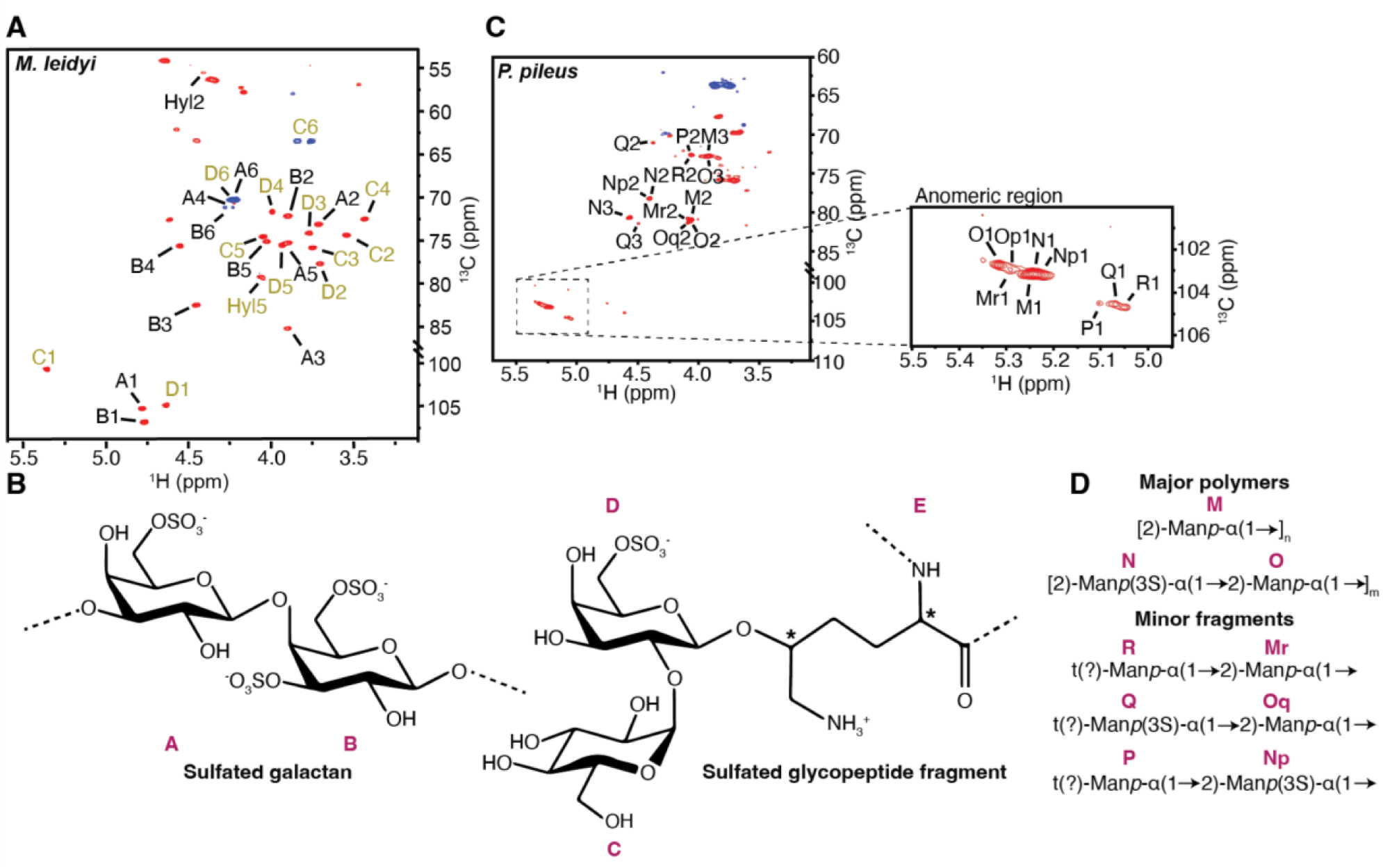
Ctenophore mesoglea contains novel sulfated polysaccharides. (**A**)^1^H-^13^C HSQC NMR spectra of an *M. leidyi* mesoglea sample, containing signals of both a sulfated galactan and a sulfated glycopeptide fragment. Anomeric and ring residue signals of the sulfated galactan disaccharide (black) and sulfated glycoprotein fragment (gold) are indicated. Unlabeled signals arose from residual peptides. (**B**) Monomer structures of the sulfated galactan polymer (residues A and B) and the sulfated disaccharide linked to hydroxylysine (residues C, D and E) found in *M. leidyi*. (**C**) ^1^H-^13^C HSQC NMR spectrum of a *P. pileus* mesoglea sample. Signals of ring C–H groups 2 and 3 are labeled in the spectrum shown on the left. On the right is an expanded anomeric region with signal assignments. (**D**) Structures of sulfated and non-sulfated mannan monomers observed in *P. pileus* with labels corresponding to the spectra.

In *P. pileus*, we found several sulfated mannan structures, as well as low levels of the same sulfated Gal-Glc-Hyl glycopeptide found in *M. leidyi*. ^1^H-^13^C HSQC NMR spectra for a *P. pileus* sample are shown in Fig. 3C with mannan structures depicted in Fig. 3D. These structures are labeled M, N, and O in the HSQC spectrum and chemical shift assignment table S3, showing different degrees of sulfation and of homo- or hetero-polymerization. Six additional minor mannose-containing residues were also identified (labeled P, Q, R, Mr, Np, and Oq), although we could resolve only a limited number of these signals, so chemical shift assignments are thus incomplete. These mannan structures are consistent with GC-MS results showing high mannose content (chromatograms shown in fig. S2B and quantified in table S2). GC-MS also showed some GalNAc and GlcNAc structures that we did not observe via NMR. These differences between techniques may result from low abundance or complex linkages to peptides. Notably, significant protein signals were observed in *P. pileus* via ^1^H NMR despite extensive protease treatment (fig. S3), suggesting the presence of complex interactions among macromolecules that are difficult to parse.

GalNAc and GlcNAc are common modifications to mucins (*41, 51, 52*). However, neither NMR nor glycoproteomics revealed protein modifications, linkages, or glycosyltransferases consistent with the glycosylation of mucins or other proteins in *M. leidyi* or *P. pileus* mesoglea. Sample digestion did produce a moderate amount of insoluble material that could not be analyzed via NMR, such that glycosylated moieties may have been missed. Similarly, O-glycosylation common to mucins can be difficult to identify without specialized sample treatments (such as mucinases), and glycosylation biochemistry is understudied (there have been no reports of glycosyltransferases in ctenophores, despite an assumption that all metazoans have glycoproteins), so more specialized studies of mucins in ctenophores are still required for a full accounting of mesoglea composition (*53*). If further studies support our preliminary finding that ctenophore mucins are unglycosylated, this may be correlated with their short PTS domains of only 20-80 residues, compared to 100+ in typical metazoan mucins (*54*). We hypothesize that ctenophore mucin proteins may instead complex with glycans via non-covalent interactions to build a mucus structure, a known property of other mucus-like materials such as biofilm complexes (*41, 55, 56*). Given the extraordinary diversity of both mucin sequences and glycans (*57*), clarifying structure-function relationships of ctenophore mucin proteins and their associated glycans will be crucial for understanding mesoglea evolution and architecture.

## Major protein constituents of ctenophore mesoglea

Via proteomics and NMR, we also observed evidence of type IV collagen. Our data further confirm the presence of only the type IV isoform, which is not known to form fibrils but instead makes a thin network in the epithelial basement membrane (Fig. 2A,B, data S1, and data S3) (*12*). The Gal-Glc-Hyl structure we identified in both species (Fig. 3B) is common in collagens, including collagen type IV (*50*), and as we did not identify any other collagen types, this structure is presumably part of ctenophores’ type IV collagen. However, total amino acid analyses indicated low relative hydroxylysine concentration in *M. leidyi*, so collagen IV is not likely to be a major constituent of the mesoglea and could come from epithelial debris (table S4). Additionally, we observed only moderate levels of glycine and proline in the mesoglea for both species; the amino acid content of collagenous tissues is typically dominated by glycine and proline, further evidence that it is unlikely that ctenophore mesoglea is primarily collagen-based (*58–60*).

Interestingly, one of the top ten protein MS hits in both *M. leidyi* and *P. pileus* had significant identity with the protein mesocentin (Fig. 2A,B, data S1, and data S3), a giant, extracellular glycoprotein for which there is only one functional description from *Caenorhabditis elegans* (*61*). Mesocentin contains multiple filamin domains interspersed with large disordered domains. In *C. elegans*, it maintains epithelial polarity and orientation of the pharynx with respect to the body wall (*61*). Given the sizable and largely acellular separation between the ctenophore epithelium and gut, it is intriguing to hypothesize similar functions for this protein in ctenophores. As is typical for MS-based proteomics, many unidentified spectra remain, so additional proteins participating in mesoglea structure remain unidentified (*62*).

## Development of a mucus skeleton

We find that ctenophore mesoglea is largely a colloidal composite of mucin and novel sulfated polysaccharides that supports sparsely distributed cells of varying types (Fig. 4A). Atomic Force Microscopy (AFM) of mesoglea showed regular 100-nm-scale colloid-like particles in the absence of an obvious network structure (figs. S4 and S5), suggesting the sulfated carbohydrates and structural proteins we documented may form colloidal aggregates in a weakly bonded colloidal gel (*40*), unlike the strong collagenous networks more typical of metazoan connective tissue. Surprisingly, synovial fluid may be a more analogous material to ctenophore mesoglea than the elastic, fibrous-collagen-based mesoglea of cnidarians. This fluid is soft yet resilient due to the presence of mucopolysaccharides and lubricin, a protein that contains mucin domains along with other non-mucin motifs (*63–65*). This finding has broad functional and evolutionary implications about the use of colloidal mucin-based materials (*66*). Mucus can have diverse functions, including physical protection, chemical filtration, and lubrication. However, animal mucus is typically secreted from the apical surface of a polarized epithelium in a thin layer external to the organism, whereas ctenophore mesoglea is a voluminous internal tissue.

**Figure 4:**
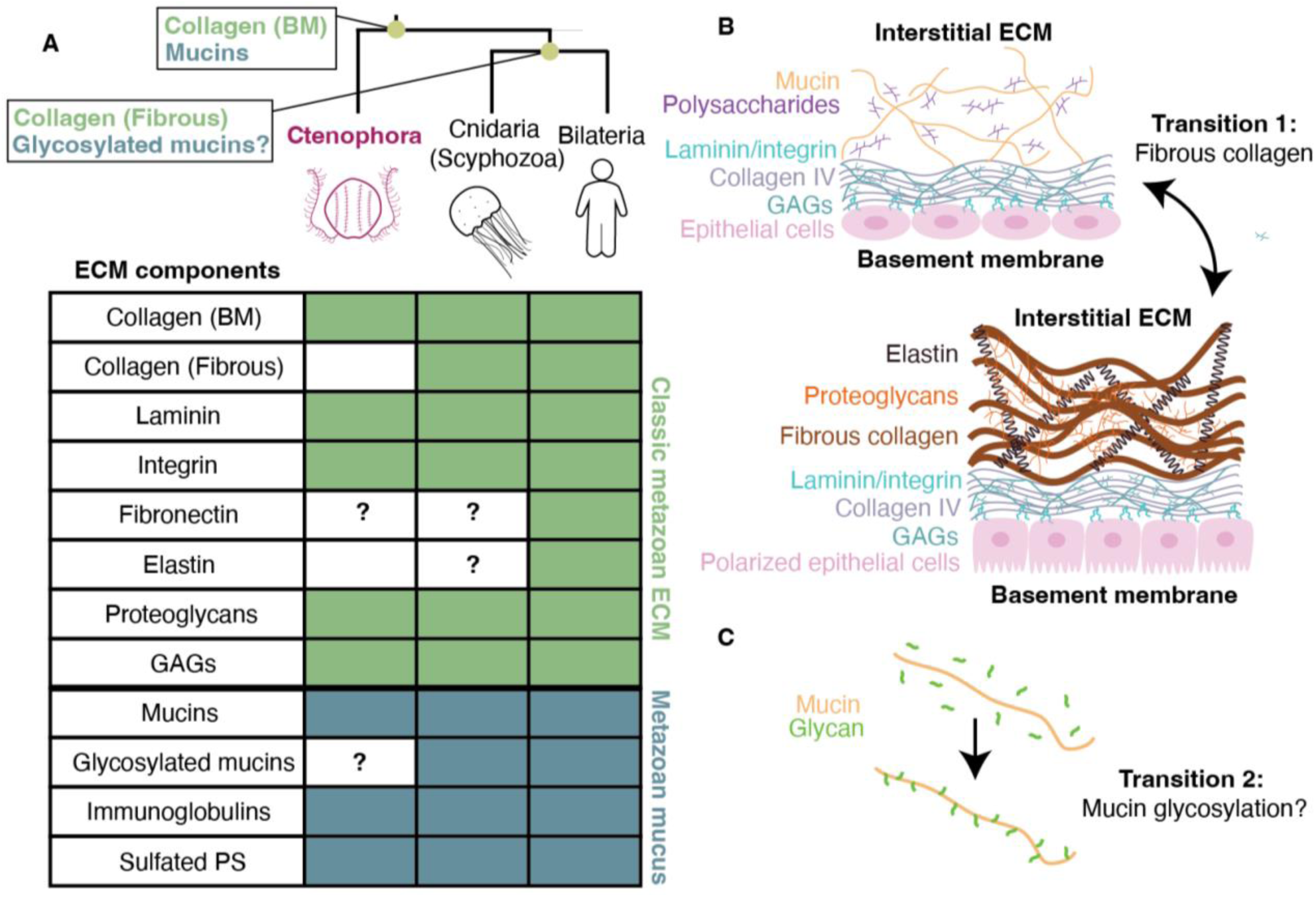
Early animal evolution shows material experimentation. **(A)** A simplified metazoan cladogram highlighting that the last common ancestor of all metazoans likely possessed mucins and basement membrane (BM) collagens. After ctenophores diverged, fibrous collagens and glycosylated mucins likely evolved, giving rise to two major extracellular material classes: the fibrous collagenous ECM (classic metazoan ECM) and mucus. Bilaterians retain the full complement of both, whereas ctenophores lack key components, leading to functional differences. Components that need further verification have a question mark. **(B)** In ctenophores, the ECM consists of mucin and polysaccharide (PS) molecules, and their epithelial cells are less organized, allowing mucus secretion into the mesoglea (top). In most metazoans, the basement membrane connects to polarized epithelial cells through integrins and laminins, creating a thin sheet of collagen type IV and glycosaminoglycans (GAGs). Their interstitial ECM is a bulk network of fibrous collagen, elastins, and proteoglycans (bottom). **(C)** A second potential transition of extracellular materials is the development of mucin glycosylation. Ctenophores do not appear to have glycosylated mucins, so their mucin proteins (orange) are likely bare and undecorated with no covalently linked glycans (green), compared to the highly decorated mucins of other metazoans.

Given its structure and composition, the nature of mesoglea development is a puzzle. In metazoans, mucus is typically produced by secretory goblet-type cells within an epithelium, and mucus-containing vesicles are released to the apical surface of the epithelium (*67, 68*). Goblet cells have been observed in ctenophore tentacles but are not well described elsewhere in the body (*55, 69–71*). Mucus is presumably produced in a similar secretory manner in ctenophores, but given the orientation of the two epithelia in the animal (the apical side of the dermal epithelium faces the external environment, and the apical side of the gut epithelium faces the interior of the gut), non-canonical secretion polarization would seem to be required for significant mucus to be deposited between the two basal surfaces of these epithelia. While unusual, alterations to epithelial polarity leading to altered geometry of mucus secretion have been documented. In certain human cancers, epithelial cell polarity inversion can occur due to defects in polarity-related gene expression, causing mucus to be secreted internally rather than externally (*72, 73*). The possibility of altered polarity of secretion is also consistent with the unusual basement membrane structure associated with epithelial tissues in some ctenophores (*12*), and with the presence of mesocentin, which alters epithelial polarity when mutated (*61,74,75*). Similarly, ctenophores lack two of nine major proteins responsible for the production and maintenance of metazoan epithelial polarity, further suggesting a less-strict regulation of the polarity of secretion in this group relative to later-branching metazoan taxa (*71, 76*).

Our results suggest two broad evolutionary findings. Firstly, while ctenophore epithelia have a basement membrane with type IV collagen, the interstitial mesoglea is a colloidal composite of mucins and novel sulfated polysaccharides and is therefore not capable of bearing loads or elastic storage of mechanical energy, in keeping with the organisms’ locomotion via macrocilia (Fig. 4B, fig. S6). Secondly, ctenophore mucin proteins appear to be minimally or possibly non-glycosylated, in sharp contrast to other metazoan mucins. This may represent a transition in animals from undecorated mucins to decorated mucins (Fig. 4C). In *M. leidyi* and *P. pileus*, the protein composition and the presence of a sulfo-glycosylated hydroxylysine associated with collagen assembly are similar. In contrast, the composition of free polysaccharides differs, perhaps reflecting different physiological requirements or other differences between the two clades within ctenophores.

In summary, we find that ctenophore mesoglea features short mucin polymers in a milieu of novel, complex polysaccharides dominated by sulfated polygalactose in *M. leidyi*, and sulfated polymannans in *P. pileus*. These macromolecules likely interact via non-covalent counterionic bonding, leading to a highly hydrated and sparse colloidal gel interspersed with actins and myosins similar to mucus in other metazoans (*36, 37*). The resulting material has a liquid-like elastic modulus (fig. S6), making it functionally distinct from the energy-storing mesoglea and connective tissues of other metazoa, in particular those of cnidarian jellyfish and sponges, whose canonical function is to store and release mechanical energy (*77–79*).

Ctenophores are among the least understood animals, and correlatedly, there is a long history of underestimating their complexity and misinterpreting their biology (*80*). Because they are round and gelatinous, they were long grouped with Cnidaria (which includes medusoid jellyfish) in a clade called Coelenterata (*9, 81*). This led to surprise when phylogenetic analyses found that they diverged from other animals earlier than cnidarians or poriferans (*3, 4*). Our findings, in the context of the more resolved placement of ctenophores as sister to other metazoans, suggest there was extensive evolutionary experimentation with material properties, macromolecular constituents, biomechanical strategies for being larger than a single cell, and the function of the clade-defining epithelium of multicellular animal bodies that occurred prior to the innovation of storage of elastic energy within a network of fibrillar collagen.

## Acknowledgments

We thank the Yale Center for Research Computing for the computational resources on the Grace cluster for BLASTp analyses, the Yale Institute for Nanoscience and Quantum Engineering for AFM usage, and Jing Yan for rheometer usage. We also thank AltaBioscience for performing the total amino acid analysis. Lauren Mellenthin, Mary Beth Decker, and other members of the Dunn Lab provided valuable assistance with ctenophore care and collection. We are also grateful to members of the Sweeney Lab, Aurelia Moriyama-Gurish, Shuhan Zhang, and Yale Peabody Museum of Natural History staff Lourdes Rojas and Eric Lazo-Wasem for additional help with ctenophore collection.

## Author contributions

Conceptualization: SS, CD AS

Methodology: SS, SAH, IB, IN, JV, PA, CD, AS

Investigation: SS, SAH, IB, IN, JV Visualization: SS, IB, JV, CD, AS Funding acquisition: SS, PA, CD, AS Supervision: PA, AS

Writing – original draft: SS, AS

Writing – review & editing: SS, JV, CD, AS

## Data, code, and materials availability

All raw data on Dryad, doi will be provided upon publication.

## Materials and Methods

### Ctenophore collection

Two species of ctenophores, *Mnemiopsis leidyi* and *Pleurobrachia pileus*, were collected via dip cups at the Connecticut shoreline of the Long Island Sound between 2020 to 2024. Ctenophores were collected under the State of Connecticut Department of Energy & Environmental Protection permits #0323011 and #0323011b. After collection, individuals were either maintained in a pseudokreisel at the salinity and temperature at which they were collected or immediately frozen at -80 ◦C.

### Mesoglea extraction

Whole individuals were vortexed for 1 minute to shear apart epithelial layers, then the whole organism was centrifuged at 8000 *g* for 15 minutes to pellet cells, nuclei, and other cellular debris. The resulting supernatant was a transparent, slightly viscous liquid. This supernatant sample was defined to be mesoglea for the purposes of these studies. This sample was decanted from the resulting pellet and used for all experiments described below. Typically, a new mesoglea sample was prepared for each experiment. A few experiments used mesoglea samples from the same individual, and these are indicated below.

### Glycosyl and glycopeptide analyses

Mesoglea samples (*M. leidyi*: n=2; *P. pileus*: n=1) were shipped to the Complex Carbohydrate Research Center at the University of Georgia for comprehensive glycosyl analyses. The mesoglea samples were preserved for two downstream experiments: (1) samples for glycoproteomics were treated with 0.02% sodium azide (ThermoFisher), 0.1 mM phenylmethylsulfonyl fluoride (PMSF, ThermoFisher), and 1X Halt Protease Inhibitor (ThermoFisher); (2) samples for all other glycosyl analyses were treated with 0.02% sodium azide and heated to 100 ◦C for 5 minutes.

Aliquots of these mesoglea samples were digested with proteases to remove proteins for carbohydrate composition experiments. First, samples underwent a double protease digestion with proteinase K (Sigma no. P4302-5G, 200 µg /10 mg sample) in 50 mM MgCl buffer (pH 7.5) overnight at 50 ◦C. This was followed by Pronase digestion (Sigma no.10165921001, 100 µg /10 mg sample) overnight at 50 ◦C. A 6 kDa MWCO dialysis membrane (Spectra Por) was used to remove digested proteins and any remaining salt from the sample. Samples were then dried using a centrifugal vacuum evaporator. These samples were used for GC-MS and nuclear magnetic resonance (NMR) experiments, described below.

GC-MS was used to perform glycosyl composition analysis, following the method outlined by Black and colleagues to produce per-O-trimethylsilyl (TMS) derivatives of the monosaccharide methyl glycosides from the digested samples (*82*). Briefly, samples are heated in 400 µl of 1 M methanolic HCl overnight at 80 ◦C and then dried with nitrogen. The samples are then re-N-acetylated by adding methanol:acetic anhydride:pyridine 2:1:1 for 20 minutes at room temperature and dried again. The sample was then derivatized with Tri-Sil® (ThermoFisher) at 80 ◦C for 30 minutes. The TMS methyl glycosides were then analyzed and visualized on the GC-MS using an Agilent 7890A GC interfaced to a 5975C MSD with a Supelco Equity-1 fused silica capillary column (30 m X 0.25 mm ID).

Dried mesoglea samples were exchanged twice from D_2_O (99.9% D, Sigma), then dried again using a centrifugal vacuum evaporator. The samples were then suspended in 520 µl of D_2_O with 50 nmol DSS-d_6_ (99.9% D, CIL), vortexed, briefly sonicated, and warmed up (∼60 ◦C), producing a slightly viscous gelatinous suspension. For *M. leidyi* samples, centrifugation was used to remove the insoluble material, and the volume of the solution was adjusted to ∼540 µl with D_2_O and transferred to a 5 mm NMR tube. For the *P. pileus* sample, the sample would not fully pellet at 16,000 *g* and was opaque. To prepare this sample for an 800 MHz spectrometer, the sample was freeze-dried and resuspended in 200 µl D_2_O (99.96% D, Sigma) and then 42 µl was transferred to a 1.7 mm NMR tube.

NMR data were collected using a Bruker Avance III at 70 ◦C with a 5 mm TCl cryoprobe (^1^H, 600.13 MHz), or a Bruker NEO spectrometer with a 1.7 mm TCl cryoprobe (^1^H, 799.17 MHz) where indicated. We used standard pulse programs in the spectrometer library to collect 1D ^1^H and 2D ^1^H, ^1^H-COSY, ^1^H, ^1^H-TOCSY, ^1^H, ^1^H-NOESY, ^1^H, ^1^H-ROESY, ^1^H-^13^C-HSQC, ^1^H-^13^C-HMBC, and ^1^H-^13^C-HSQC-TOCSY (*M. leidyi* only) experiments. A total recycling delay of 60 seconds was used while collecting quantitative ^1^H NMR data. We used a mixing time of 60 to 120 ms for NOESY and 200 ms for ROESY experiments. The spin-lock time was 60 or 120 ms for TOCSY and 70 ms for HSQC-TOCSY. NMR data were processed in Topspin 4.0.5 (Bruker) and analyzed in CCPN Analysis 2.4 (*83*) or 3.2.0 (*84*). We rendered figures in MestReNova 14.2.3 (Mesterelab Research). We referenced ^1^H and ^13^C chemical shifts to the respective DSS signals at 0.00 ppm.

An aliquot of one of the digested *M. leidyi* mesoglea samples underwent anion exchange chromatography to further investigate the presence of potentially sulfated polysaccharides. A 5 ml DEAE HiTrap column (GE) was used to perform high-performance liquid chromatography analysis (HPLC). 250 µl of a 5 mg /ml sample (dissolved in water) was injected first and run for four minutes with 100% water. Then, the buffer was switched to a 2 M NH_4_OAc solution over 31 minutes and held at 100% 2 M NH_4_OAc for five minutes. 100% water was then re-injected into the column until equilibration for the last minute of the run. Sample fractions were collected and dried at one-minute intervals and reconstituted in 200 µl of DI water. Sulfated galactans were not observed after fractionation because they only eluted with high NaCl concentration.

Verification of carbohydrate presence in the fractions was detected using a phenol sulfuric acid assay. We used a modified version of the microscale phenol sulfuric acid assay of Masuko and colleagues (*85*). In short, 0.2 ml of concentrated sulfuric acid was added to wells containing 10 µl of resuspended sample fractions. 5% phenol was added to the wells, and plates were incubated at 50 ^◦^C for 20 minutes. Absorbance was read at 490 nm using a Molecular Devices Spectra Max plate reader. Fraction four of the DEAE separation contained carbohydrates and was then run on the NMR according to the method above.

### Glycoproteomics

All of the following reagents were purchased from Sigma Aldrich unless otherwise mentioned. Mass spectrometric data was performed on a Thermo Scientific LTQ Orbitrap Eclipse Tribid mass spectrometer attached with an Ultimate RSLCnano-LC system. Data analysis was performed using Xcalibur 3.0 and Byonic v4.10 software.

Proteomics of both tryptic digests from one sample of *M. leidyi* and one *P. pileus* sample, as well as their pre-digested samples, were performed using LC-MS/MS (Liquid Chromatography-MS/MS). The provided samples (pre-digested or not) were reduced, alkylated with iodoacetamide, and the protein was digested with (1:10) sequencing-grade trypsin (Promega) at 37 ◦C for 16 hours. The peptides were analyzed on an Orbitrap Eclipse or Orbitrap Ascend mass spectrometer equipped with a nanospray ion source. Pre-packed Nano-LC columns of 15 cm length with 75 µm internal diameter, filled with 3 µm C18 reverse-phase material, were used for chromatographic separation of the samples. The separation conditions were low to high acetonitrile in a solution containing 0.1% formic acid, and the separation time was 60 minutes. The precursor ion scan was acquired at 120,000 resolution in an Orbitrap analyzer, and precursors at a time frame of 3 seconds were selected for subsequent fragmentation using higher-energy collisional dissociation (HCD). The threshold for triggering an MS/MS event on the ion-trap was set to 500 counts. Charge state screening was enabled, and precursors with an unknown charge state or a charge state of +1 were excluded. Dynamic exclusion was enabled (exclusion duration of 30 s). The fragment ions were analyzed on an Orbitrap of HCD at 30,000 resolution.

The resulting data were either hand-processed or processed with Byonic and searched against protein models for each species (described below) and a catalog of basic N-glycans and O-glycans. Post-translational modifications, including oxidation and deamidation, were included in the variable search. Carboxyamidomethylation of the cysteine was included as a fixed modification. The precursor mass tolerance and fragment mass tolerances were set to 10 ppm and 20 ppm, respectively.

For *M. leidyi*, we used protein models from the recently published chromosome-level genome for the northern Atlantic *M. leidyi* population, accessed on GenBank under GCA 963919725.1 (*26*). For *P. pileus*, there are no genome or assembled protein models available so we used protein models from the closely related *Hormiphora californensis* chromosome-level genome, generated by Schultz et al. (2023) and accessible on GenBank under GCA 020137815.1 (*4*). We provided the functional annotation of each identified protein and glycoprotein through sequence and structural searches. For sequence searches, BLASTp, Pfam, and Gene Ontology terms were queried for both species. For both species, we used BLASTp with the NCBI nr database (April 2025 for *M. leidyi,* June 2025 for *H. californensis*) with an e-value cut-off of 1E-5 (*86*). For further functional annotation, we used InterProScan via its webpage (https://www.ebi.ac.uk/interpro/search/sequence/) to identify protein family domains and GO terms (*28*). We used the default parameters with all databases available. The MOTIF Search tool on https://www.genome.jp/ was also used for domain architecture visualization.

Several proteins for both species did not contain any results via the sequence searches or only had identity to hypothetical proteins with no functional annotations, so structural searches were also performed for those particular proteins. Additionally, structural searches were conducted for proteins of particular interest, such as collagens and mucins, described further in this paper. We used AlphaFold3 to predict structures via the online server: https://alphafoldserver.com/ (*87*). We searched for any homology and thus used structure predictions regardless of pLDDT accuracy to maximize the likelihood of finding any small structural similarities. We then used the top-ranked prediction for each protein and searched for structural similarity to any other proteins in the PDB or AlphaFold DB via FoldSeek at https://search.foldseek.com/search (*88*). The e-values of all hits with structural homology are reported in our supplementary data (data S2 and S4). Any given protein’s particular structural details would need to be further investigated with confirmed high-confidence structure predictions to show true structural homology.

For proteins with potential mucin homology, we used PTSPred, a Perl-based PTS/mucin domain search tool developed by Lang and colleagues (2004), to identify PTS domains (*29*). For some mucin-like proteins, we also performed visual inspections to identify PTS-rich sequences that were too short to be detected with the search tool. Cysteine knots were also identified using the motif search on ScanProsite with the consensus motif C-x-G-x-C (*89, 90*). Illustrator for Biological Sequences 2.0 was used for protein domain architecture visualization (*91*).

### Total amino acid analysis

Two biological replicates of *M. leidyi* and *P. pileus* mesoglea were sent to a contract service to determine their total amino acid composition (AltaBiosciences, Redditch, UK). Upon arrival, the mesoglea samples were degassed, sealed under vacuum, and hydrolyzed for 24 hours in 6 M HCl. After vacuum hydrolysis, the samples were dried to remove the concentrated HCl and diluted to the required volume using 0.1 M HCl buffer. The hydrolyzed samples were analyzed via ion-exchange chromatography with post-column ninhydrin detection for quantification of each amino acid (Biochrom 30+, Harvard Biosciences, Holliston MA). Hydroxyproline and hydroxylysine were separately identified using absorbance at 440 nm and 570 nm, respectively.

### Atomic Force Microscopy (AFM)

We used 15 mm muscovite mica discs (Electron Microscopy Sciences, P/N: 71856-03) as a surface for sample testing, freshly cleaving a mica slice for each sample. We then deposited 5 µl of each mesoglea sample onto the mica disc. We tried several sample layering techniques to identify the most successful protocol, including wet samples, simple air dry, and using a spin coater to level and thin the sample and then air drying. The most successful protocol was depositing a sample onto the disc then immediately applying the disc to a spin coater. After a few minutes of spinning, the disc was removed, producing a thin layer of dried sample. Other sample preparation methods did not produce clear images. We used a few mesoglea samples from different specimens of *M. leidyi* and *P. pileus* for testing and verified the protocol using *𝜅-*carrageenan, successfully visualizing the gel network.

A Bruker Dimension FastScan Atomic Force Microscope was used to understand the structural morphology of ctenophore mesoglea. Mica discs with prepared samples were placed onto the granite platform to begin scanning. We tried multiple tip types and found SCANASYST-AIR and FASTSCAN-B tips (Bruker) to pair best with our soft mesoglea samples. For each experiment, a new tip was mounted onto the cantilever. The AFM was used in Scan Assist Mode (peak force mode) with the Scan Assist in Air setting. The laser was calibrated and aligned at the start of each experiment. The following parameters were used for each experiment: starting scan size of 15 µm, aspect ratio of 1, no x/y offset, scan rate of 4 Hz or 1 Hz, samples/line at 256 or 512, ScanAsyst Auto Control on, peak force amplitude at 150 nm, and lift height of 300 nm. Each scan produced a Height Sensor Image and a Peak Force Error Image. For each experiment, clear images were saved and exported for interpretation.

### Rheology

All rheology experiments were performed on a shear rheometer (Anton Paar Physica MCR502WESP) in strain-control modes. Various measuring geometries, such as cone-and-plate and parallel plate, were initially used, but yielded noisy results. Therefore, a double gap system was used to maximize contact area. The inner and outer cylinders sandwich the rotational cup, which provides a large surface area for the sample to contact the measuring system. The double gap system used was compatible with the MCR equipment (Anton Paar DIN 54453; measuring cylinder: B-DG26.7Q1, measuring cup C-DG26.7T200SS). A Peltier Temperature Control Device (Anton Paar C-PTD200) designed for the double gap geometry was used to maintain constant temperature during all experiments. The temperature was set at 17 ◦C, consistent with the temperature of the ctenophore habitat.

Only *M. leidyi* results are reported here due to low sample size of *P. pileus* being available at the time of rheometry experiments. Approximately 5 ml of ctenophore mesoglea was deposited via pipette between the inner and outer cylinders. Controls of soybean oil, gelatin, and water were used to ensure the system was operating properly.

We used amplitude sweeps to identify the linear viscoelastic strain range, then we measured the viscoelastic properties, specifically elastic and viscous moduli, using frequency sweeps. Mesoglea from a total of three *M. leidyi* specimens were used for amplitude sweeps. Here, we reported elastic and viscous moduli extracted from the viscoelastic region of frequency sweeps, specifically values resulting from shear strain applied at 0.01 Hz and 1% strain. All relevant amplitude and frequency sweep experiments are included in the raw data.

In brief, the elastic and viscous moduli are calculated as follows. Upon applying shear strain, the rheometer recorded the time (*seconds*), deflection (*radians*), torque (*N-m*), applied shear strain (*%*), and shear stress response (*Pa*). For this study, shear strain (Eq. S1) is defined as *γ* (unitless), shear deformation, *s* (*meters*), and shear gap, *h* (*meters*). Since we are using double gap geometry, *ℎ* is a fixed value (often defined as an instrument-specific factor) and *s* is equal to the radius of the cylinder at the liquid interface *𝑟* times the angular displacement, *θ*, in radians.

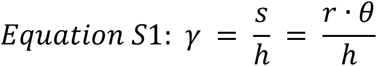

Shear stress is defined by Equation S2 with *τ* as shear stress, *F* for force (*N*), and A for area

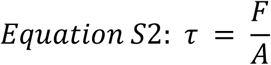

A basic shear modulus can be calculated to describe a material’s properties at specific time points (Equation S3).

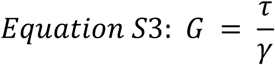

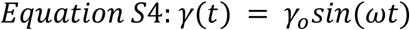

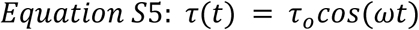

The strain is applied as an oscillatory shear by the rheometer and the stress response results in a waveform, noted in Equations S4 and S5, where *γ_o_* is the strain amplitude, *τ_o_* is the stress amplitude, *ω* is the angular frequency, and *δ* is the phase lag between the stress and strain waveforms. To ensure that the signal of the stress response is higher than instrument noise, we evaluated the stress response waveforms in relation to the applied strain. If a given strain produced a sinusoidal stress response, the strain was considered valid, and Microsoft Excel was used to record a database of (elastic) modulus, *G’*, and loss (viscous) modulus, *G”*, from the RheoCompass software as a function of *γ*. *G’* is defined in Equation S6 and *G”* is defined in Equation S7.

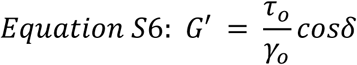

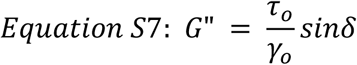

The viscoelastic moduli were visualized as a function of *γ* for the valid strain range using MATLAB 2023a. The linear viscoelastic (LVE) region was evaluated via visual inspection by identifying regions where *G’* and *G”* exhibit plateaus across strains. The LVE region is defined as the strain amplitude range where viscoelastic moduli are independent of strain. Once we identified the strain values that produced viscoelastic moduli within the LVE region, we performed frequency sweeps. A frequency range, such as 0.01-5 Hz, starting with the highest frequency and decreasing to the slowest frequency until the sweep was complete. We then performed the same analysis described above for the amplitude sweeps to evaluate the raw waveform data and ensure we only continued studying samples with high signal and low noise.

Then, we created a database of the frequencies that produced data with a sinusoidal stress response. Using the understanding of the LVE region, we extracted the calculated values of the storage (elastic) modulus, *G’*, and loss (viscous) modulus, *G”*, from the RheoCompass software as a function of *𝜈* for sweeps performed with strains in the LVE region, such as 1%. The viscoelastic moduli were visualized compared to other tissues using MATLAB 2023a.

**Fig. S1.**
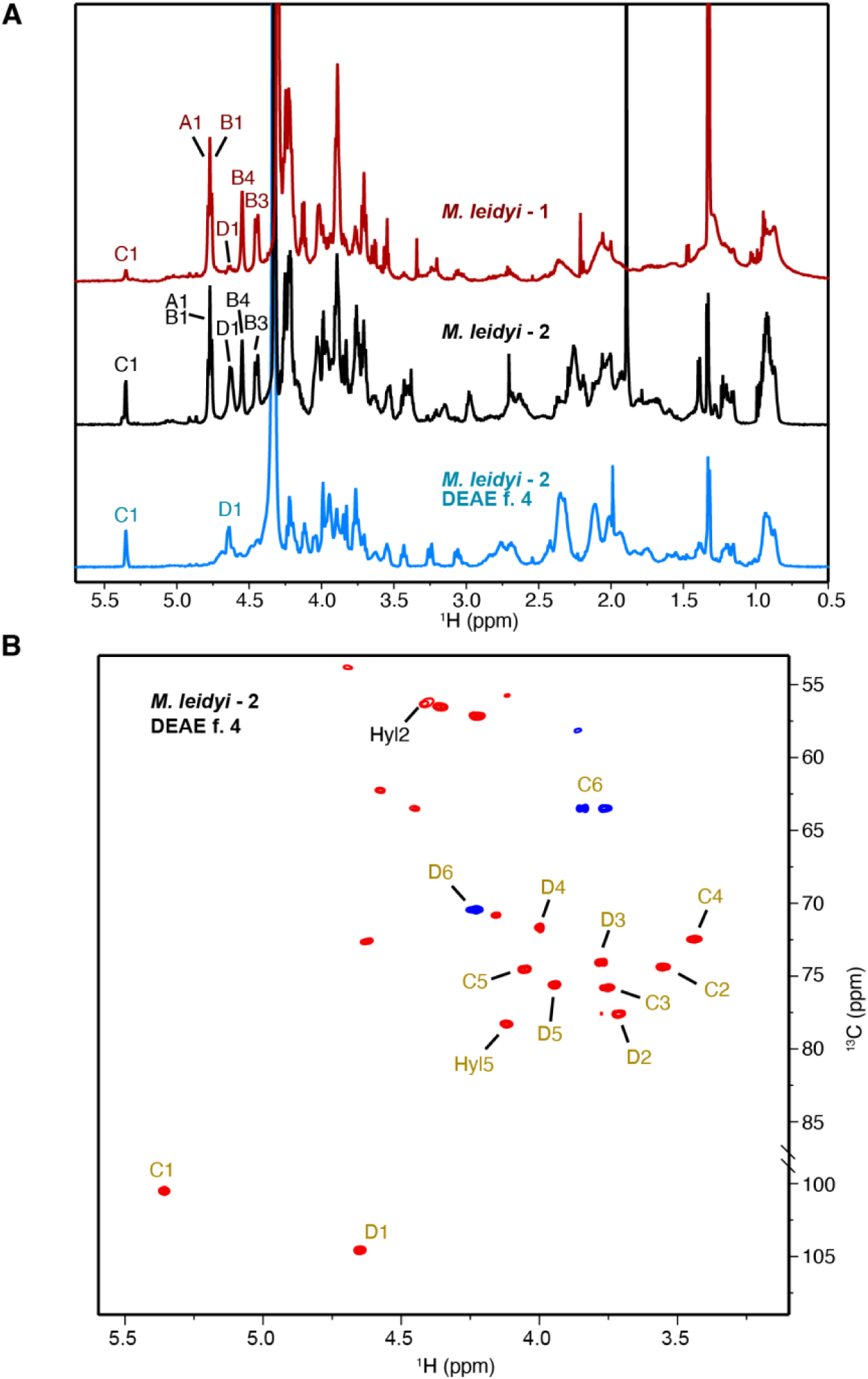
Unique sulfated galactans are present in *M. leidyi* mesoglea. (A) ^1^H NMR spectra of two *M. leidyi* mesoglea samples. The bottom sample is DEAE fraction 4 of the previous *M. leidyi* sample. (B) ^1^H-^13^C HSQC NMR spectra of the DEAE fraction 4 of the second *M. leidyi* mesoglea sample, where only signals of sulfated glycopeptide fragment were present, confirming that sulfated galactan and sulfated glycoprotein were not covalently bound. The sulfated glycan fragment (gold) and the amino acid fragment (black) are indicated. Unlabeled signals arose from residual peptides.

**Fig. S2.**
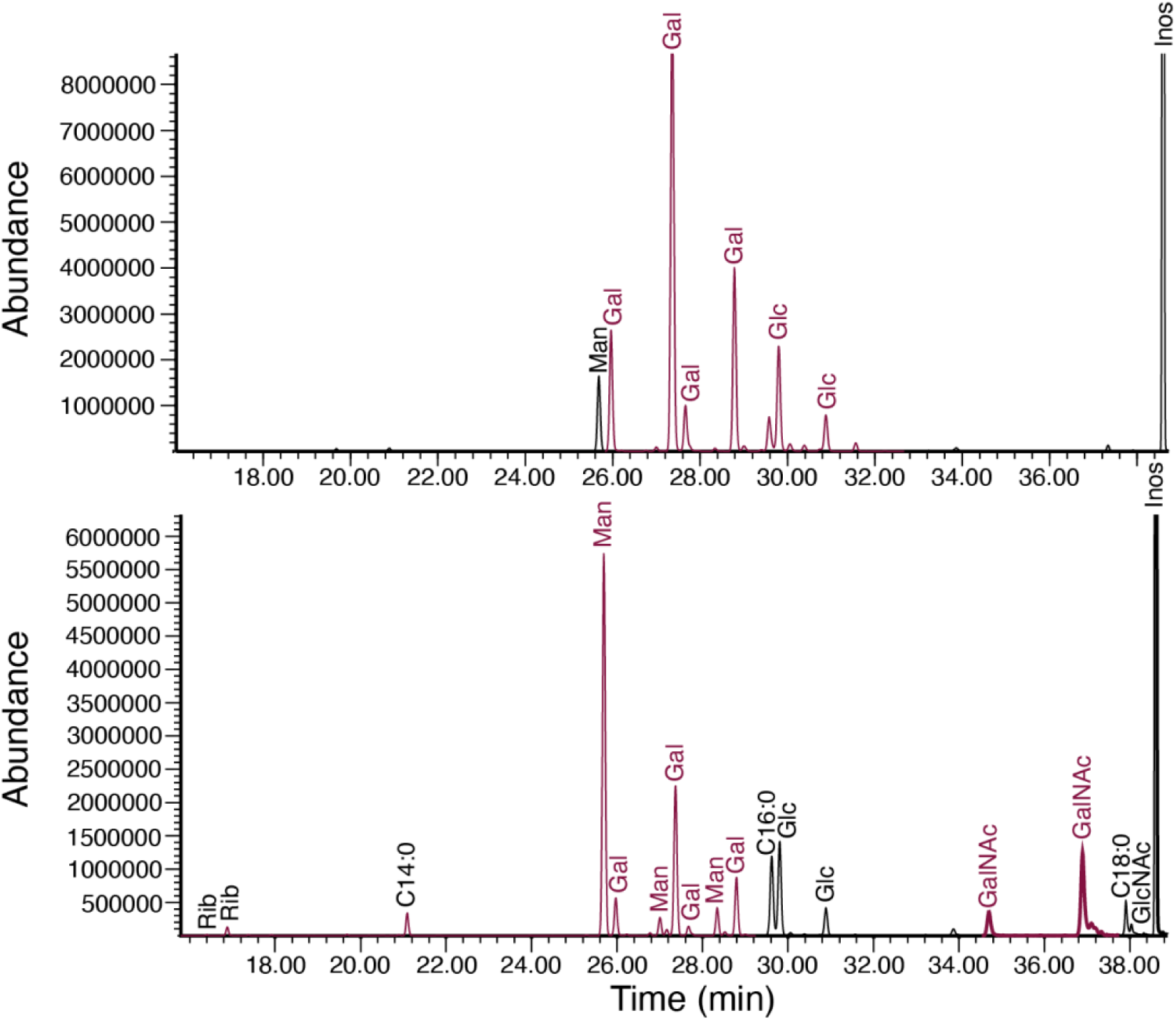
Galactose-based monosaccharides are abundant in ctenophore mesoglea. (A) Total ion chromatograms from GC-MS of *M. leidyi*, highlighting high galactose content. (B) Total ion chromatograms from GC-MS of *P. pileus*, highlighting high mannose content. Most abundant peaks are highlighted in red. Chromatograms were taken after derivatization of the sample that underwent protease treatment and dialysis. n=1 for both species, and inositol is present in both plots as a reference.

**Fig. S3.**
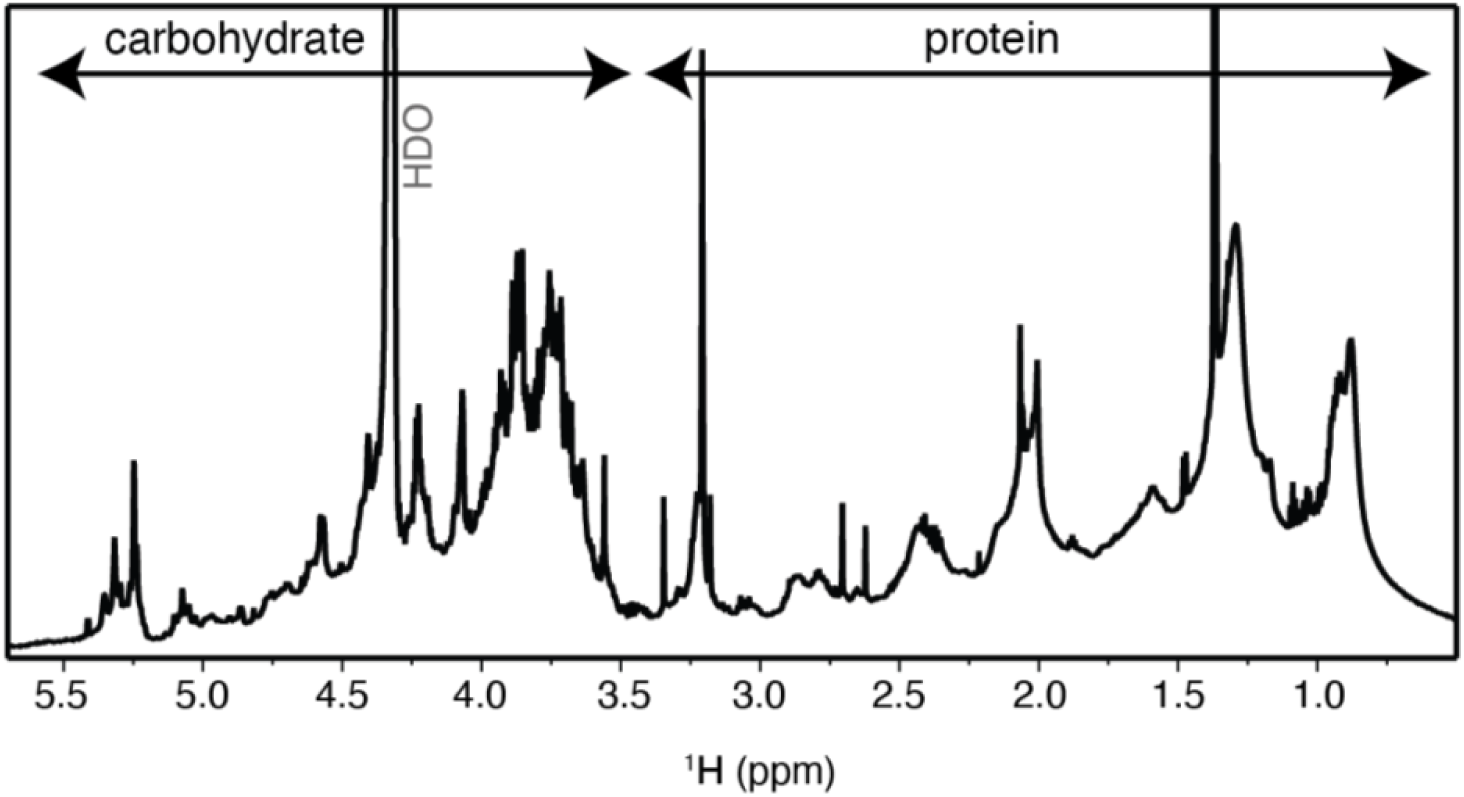
Sulfated mannan structures and residual protein found in *P. pileus*. ^1^H NMR spectra of one *P. pileus* mesoglea sample, highlighting significant protein and carbohydrate signal.

**Fig. S4:**
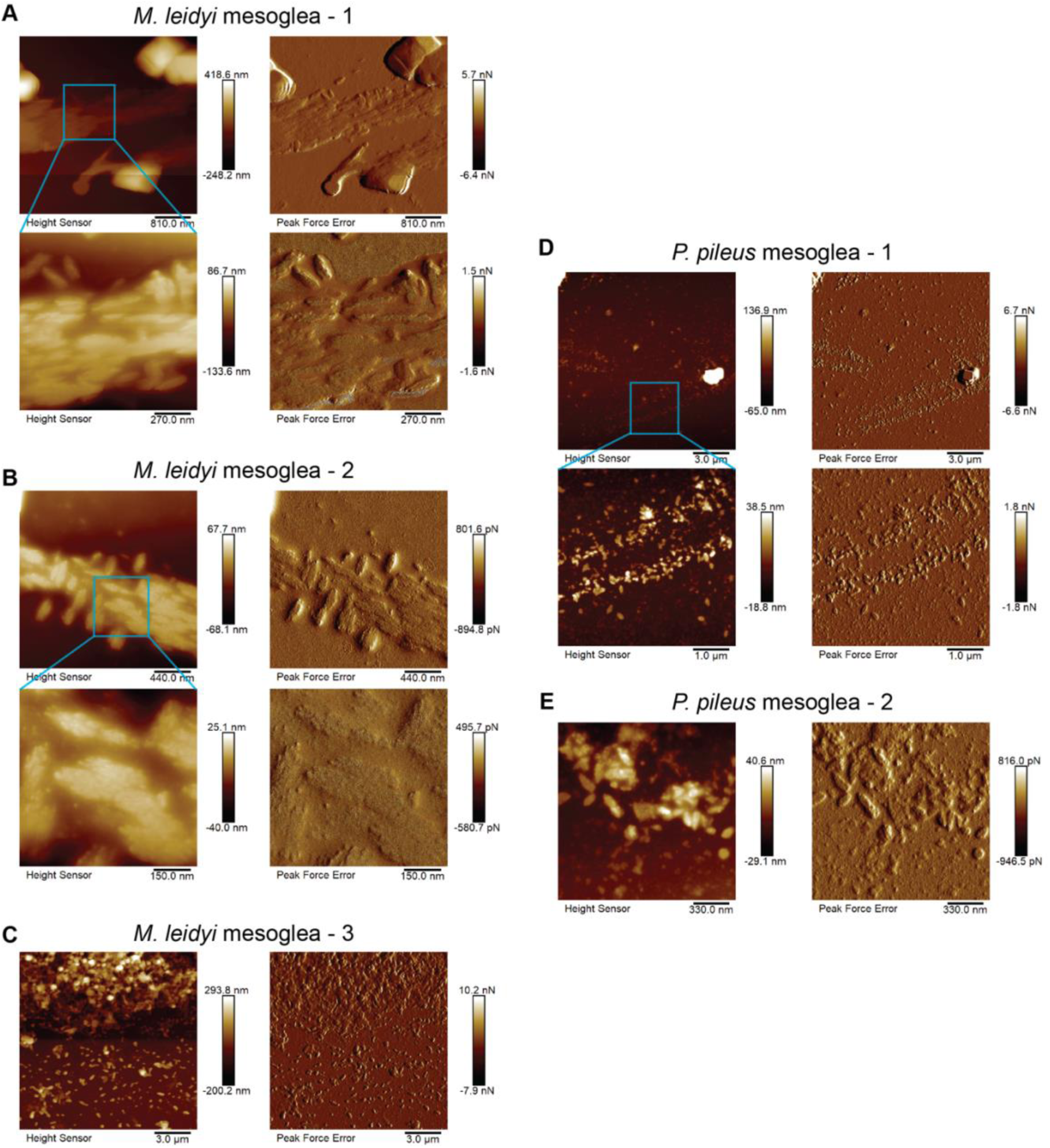
AFM height sensor and peak force error images representing organized colloidal structures in ctenophore mesoglea. (A): An example of *M. leidyi* mesoglea, zooming in on an organized structure featuring rod-like morphology at 800 nm and 270 nm. The left image in all AFM results is the height sensor image and the right image is the peak force error image. (B): Another *M. leidyi* mesoglea sample showing organized structures with rod-like morphology at 400 nm and 150 nm. (C): An additional *M. leidyi* mesoglea sample showing a dense, organized structure, with many aggregates in a gradient at 3 µm. (D): A representative sample of *P. pileus* mesoglea showing two aggregated, organized structures at two levels, 3 µm and 1 µm, both also highlighting a speckled background. (E): Another P. pileus sample showing an aggregate of rod-like structures with speckled background at 330 nm.

**Fig. S5:**
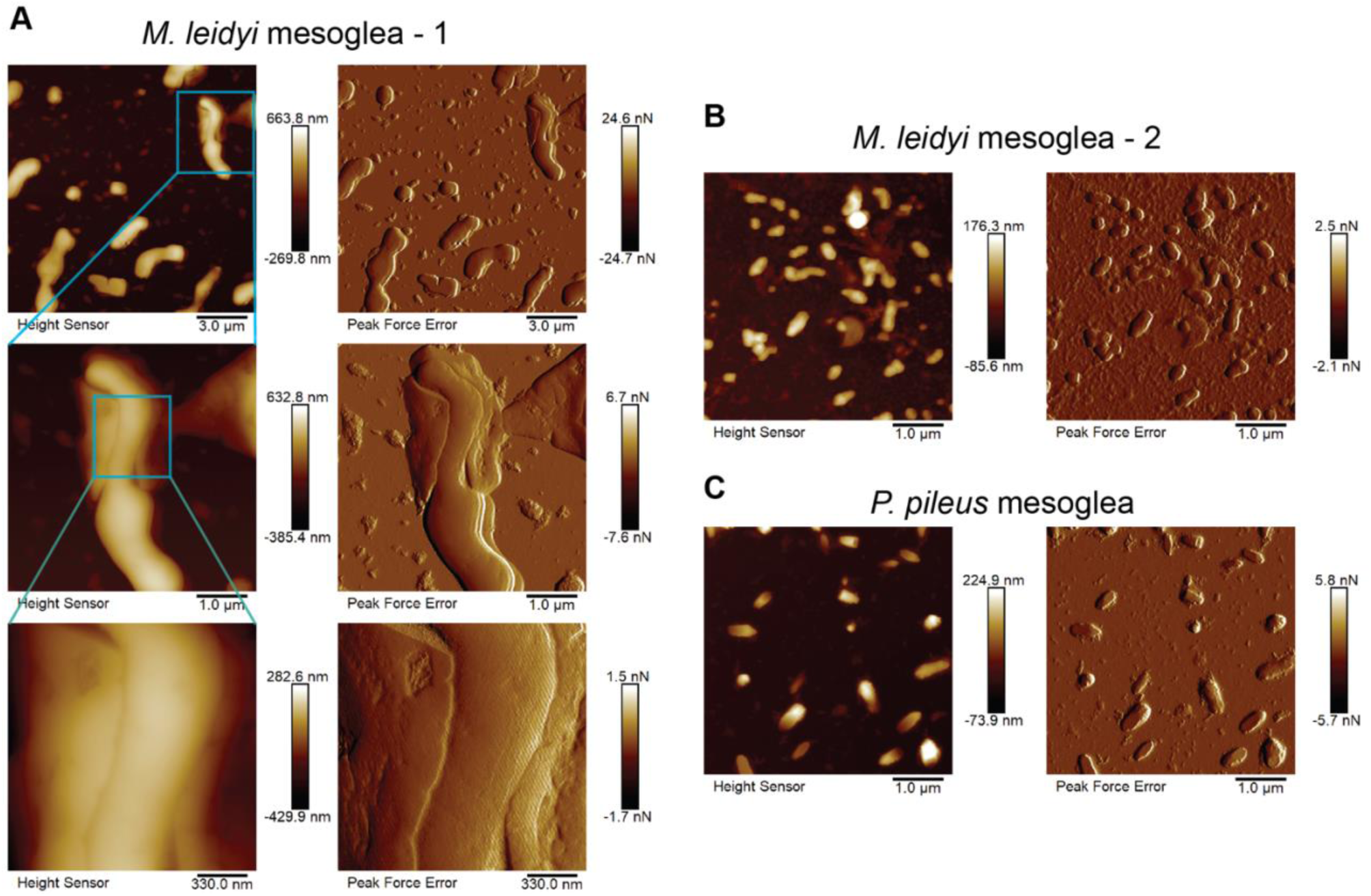
AFM height sensor and peak force error images representing amorphous blob colloidal structures in ctenophore mesoglea. (A): An example of *M. leidyi* mesoglea, zooming in on a particular aggregate structure. (B): Another *M. leidyi* mesoglea sample showing a variety of blob structures at 1 µm. (C): A representative *P. pileus* mesoglea sample exhibiting blob structure at 1 µm.

**Figure S6:**
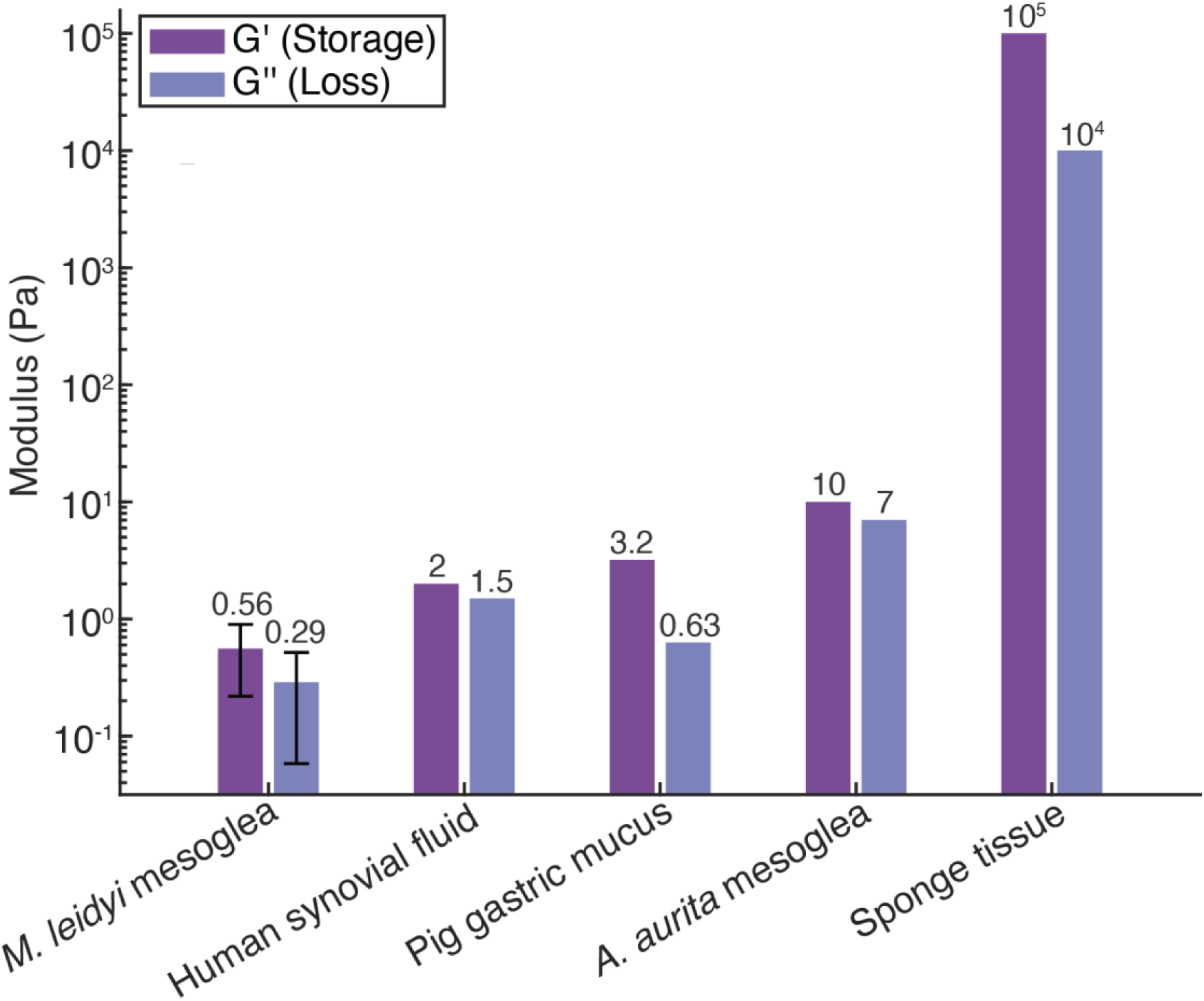
Rheological properties of ctenophore mesoglea are more similar to fluids than typical viscoelastic tissues in metazoans. Average viscoelastic moduli of *M. leidyi* mesoglea compared to other metazoan tissues (*63, 78, 79, 92*). *G’* and *G”* for *M. leidyi* mesoglea are extracted from the viscoelastic region of a frequency sweep and represent viscoelastic properties from shearing applied at 0.01 Hz and 1% strain. n=3 for *M. leidyi* mesoglea and error bars represent standard error.

**Table S1.** ^1^H-^13^C NMR chemical shifts of galactan residues found in *M. leidyi* (Fig. 3A,B and fig. S1).

| ID | Residue | Chemical shift (ppm) |  |  |  |  |  | Links |
| --- | --- | --- | --- | --- | --- | --- | --- | --- |
|  |  | 1 | 2 | 3(a,b) | 4 | 5 | 6(a,b) |  |
| A | $\rightarrow 3$)-Galp(6S)-β(1→ | 4.78<br>105.4 | 3.71<br>73.5 | 3.90<br>85.4 | 4.25<br>71.6 | 3.90<br>75.6 | **4.23**<br>**70.6** | A1→B4 |
| B | $\rightarrow 4$)-Galp(3S,6S)-β(1→ | 4.77<br>107.0 | 3.90<br>72.5 | **4.45**<br>**82.8** | 4.56<br>76.0 | 4.03<br>75.5 | **4.26**<br>**71.5** | B1→A3 |
| C | t-Glcp-α(1→ | 5.36<br>100.5 | 3.55<br>74.4 | 3.75<br>75.8 | 3.44<br>72.5 | 4.05<br>74.5 | 3.84, 3.76<br>63.5 | C1→D2 |
| D | $\rightarrow 2$)-Galp(6S)-β(1→ | 4.65<br>104.6 | 3.71<br>77.6 | 3.78<br>74.0 | 4.00<br>71.7 | 3.95<br>75.6 | **4.23**<br>**70.4** | D1→Hyl5 |
| E | Hyl | 176.2 | 4.36<br>56.5 | 2.02, 1.84<br>29.1 | 1.75<br>31.2 | 4.12<br>78.3 | 3.25, 3.07<br>45.5 |  |

*Italicized values* indicate glycosyl linkage positions; **bold values** indicate sulfation.

**Table S2.**
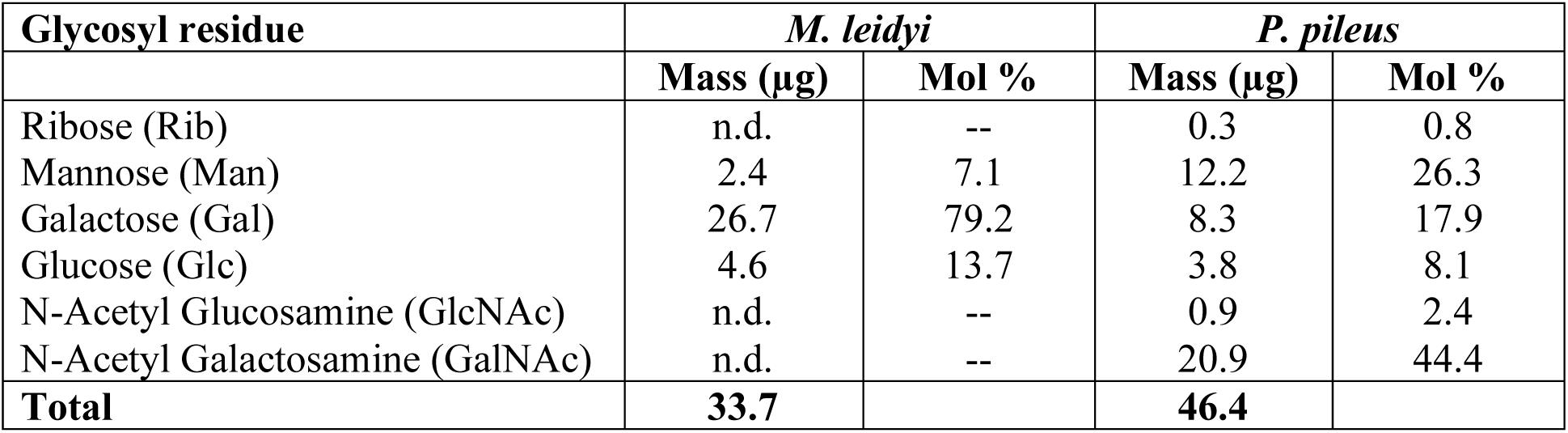
GC-MS (fig. S2) highlights high galactose in ctenophore mesoglea. (n.d. = no data)

**Table S3.**
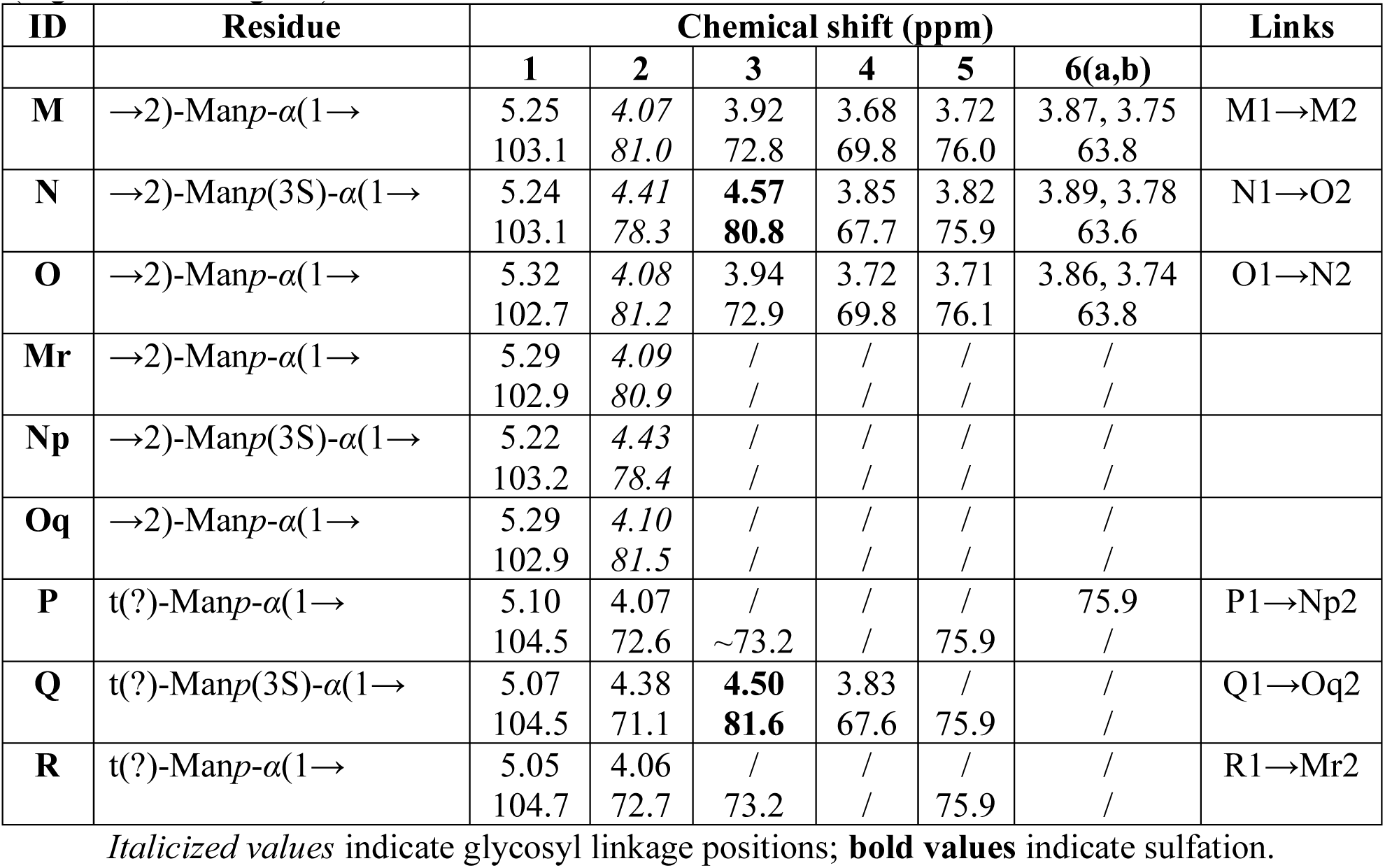
^1^H-^13^C NMR chemical shifts of sulfated mannan residues found in *P. pileus*

| ID | Residue | Chemical shift (ppm) |  |  |  |  |  | Links |
| --- | --- | --- | --- | --- | --- | --- | --- | --- |
|  |  | 1 | 2 | 3 | 4 | 5 | 6(a,b) |  |
| **M** | $\rightarrow 2$)-Manp-α(1→ | 5.25<br>103.1 | 4.07<br>*81.0* | 3.92<br>72.8 | 3.68<br>69.8 | 3.72<br>76.0 | 3.87, 3.75<br>63.8 | M1→M2 |
| **N** | $\rightarrow 2$)-Manp(3S)-α(1→ | 5.24<br>103.1 | 4.41<br>*78.3* | **4.57**<br>**80.8** | 3.85<br>67.7 | 3.82<br>75.9 | 3.89, 3.78<br>63.6 | N1→O2 |
| **O** | $\rightarrow 2$)-Manp-α(1→ | 5.32<br>102.7 | 4.08<br>*81.2* | 3.94<br>72.9 | 3.72<br>69.8 | 3.71<br>76.1 | 3.86, 3.74<br>63.8 | O1→N2 |
| **Mr** | $\rightarrow 2$)-Manp-α(1→ | 5.29<br>102.9 | 4.09<br>*80.9* | /<br>/ | /<br>/ | /<br>/ | /<br>/ | |
| **Np** | $\rightarrow 2$)-Manp(3S)-α(1→ | 5.22<br>103.2 | 4.43<br>*78.4* | /<br>/ | /<br>/ | /<br>/ | /<br>/ | |
| **Oq** | $\rightarrow 2$)-Manp-α(1→ | 5.29<br>102.9 | 4.10<br>*81.5* | /<br>/ | /<br>/ | /<br>/ | /<br>/ | |
| **P** | t(?)-Manp-α(1→ | 5.10<br>104.5 | 4.07<br>72.6 | /<br>*~73.2* | /<br>/ | /<br>75.9 | 75.9<br>/ | P1→Np2 |
| **Q** | t(?)-Manp(3S)-α(1→ | 5.07<br>104.5 | 4.38<br>71.1 | **4.50**<br>**81.6** | 3.83<br>67.6 | /<br>75.9 | /<br>/ | Q1→Oq2 |
| **R** | t(?)-Manp-α(1→ | 5.05<br>104.7 | 4.06<br>72.7 | /<br>73.2 | /<br>/ | /<br>75.9 | /<br>/ | R1→Mr2 |

*Italicized values* indicate glycosyl linkage positions; **bold values** indicate sulfation.

**Table S4.** Ctenophore mesoglea is rich in negatively charged amino acids.

| **Amino Acid** | ***M. leidyi***<br>**(µg/ml)** | **% total** | ***P. pileus***<br>**(µg/ml)** | **% total** |
| --- | --- | --- | --- | --- |
| Glutamic acid | 86 ± 18 | 18.9 ± 0.3 | 333 ± 63 | 14.2 ± 0.0 |
| Aspartic acid | 55.8 ± 10.7 | 12.4 ± 0.0 | 234 ± 28 | 10.1 ± 0.7 |
| Arginine | 24.5 ± 4.4 | 5.44 ± 0.07 | 217 ± 55 | 9.12 ± 0.59 |
| Lysine | 19.8 ± 3.3 | 4.42 ± 0.11 | 173 ± 30 | 7.38 ± 0.14 |
| Leucine | 19.2 ± 3.6 | 4.26 ± 0.01 | 166 ± 41 | 6.98 ± 0.40 |
| Valine | 24.1 ± 4.3 | 5.35 ± 0.07 | 150 ± 35 | 6.34 ± 0.29 |
| Glycine | 45.8 ± 10.2 | 10.1 ± 0.3 | 131 ± 12 | 5.69 ± 0.57 |
| Threonine | 24.4 ± 4.7 | 5.41 ± 0.02 | 126 ± 31 | 5.33 ± 0.29 |
| Isoleucine | 15.3 ± 3.1 | 3.38 ± 0.04 | 114 ± 26 | 4.83 ± 0.19 |
| Phenylalanine | 13.7 ± 2.6 | 3.03 ± 0.01 | 112 ± 24 | 4.75 ± 0.12 |
| Proline | 31.2 ± 4.4 | 6.99 ± 0.35 | 110 ± 12 | 4.78 ± 0.41 |
| Alanine | 22.5 ± 4.8 | 4.97 ± 0.12 | 102 ± 11 | 4.40 ± 0.35 |
| Tyrosine | 7.83 ± 0.45 | 1.82 ± 0.44 | 101 ± 27 | 4.25 ± 0.33 |
| Serine | 16.2 ± 2.5 | 3.62 ± 0.13 | 95 ± 17 | 4.05 ± 0.03 |
| Methionine | 7.31 ± 1.30 | 1.63 ± 0.02 | 58.1 ± 11.8 | 2.47 ± 0.03 |
| Histidine | 6.43 ± 1.12 | 1.43 ± 0.02 | 47.9 ± 8.5 | 2.05 ± 0.03 |
| Cystine | 9.7 ± 1.4 | 2.16 ± 0.09 | 47.8 ± 12.7 | 2.01 ± 0.16 |
| Hydroxyproline | 9.1 ± 3.3 | 1.95 ± 0.36 | 13.0 ± 2.9 | 0.549 ± 0.017 |
| Hydroxylysine* | 12.6 ± 3.0 | 2.78 ± 0.13 | 11.9 ± 0.4 | 0.527 ± 0.115 |
| **Totals** | **451 ± 86** | **100%** | **2345 ± 445** | **100%** |

n=2 per species, ± SE. \*Percentage recovery not fully validated.

**Data S1. (separate file) BLASTp annotations of *M. leidyi* proteomics results highlight many structural and ECM proteins.** Each MS proteomics hit that had identity with a known ECM or load-bearing protein is listed. For each protein, annotations of three major BLAST hit are included with accession numbers, e-values, and identity percentages. For the protein, the log probability from the LC-MS/MS results is also included.

Data S2. (separate file) FoldSeek annotations of *M. leidyi* proteomics results highlight additional structural and ECM proteins. For the proteins that did not result in any functional information from BLAST searches, AlphaFold and FoldSeek were used to search for structural homology to any known proteins. Through this method, we identified a couple of additional proteins that may play roles in the ECM. For each protein, the top FoldSeek hit is listed with its accession number, e-value, identity percentage, the FoldSeek database that was used, and the log probability from the proteomics results.

**Data S3. (separate file) BLASTp annotations of *P. pileus* proteomics results highlight many structural and ECM proteins.** The results for the *P. pileus* sample are listed in the same format as data S1, using predicted proteins from the *H.californensis* genome.

Data S4. (separate file) FoldSeek annotations of *P. pileus* proteomics results highlight additional structural and ECM proteins. The results for the *P. pileus* sample are listed in the same format as data S2, using predicted proteins from the *H.californensis* genome.

